# Single virus lipid mixing study of Sendai virus provides insight into fusion mechanism

**DOI:** 10.1101/2025.07.23.666410

**Authors:** Lisa Ji, Daniel Yuan, Abraham Park, Katherine Bai, Robert J. Rawle

**Author notes:** **<u>Correspondence</u>**.

## Abstract

Single virus studies have proven useful to interrogate the entry mechanism for several viral families. Here, we employ a fluorescence microscopy-based single virus assay to study the fusion (lipid mixing) of Sendai virus to model membranes, the first for any paramyxovirus to our knowledge. We find that fusion wait times following binding are exponentially distributed, suggesting a single rate-limiting step. Compared to previously studied viruses, fusion is relatively slow (tens of minutes) and inefficient (only a small fraction of virions undergo fusion). Trypsin treatment of the virus or different viral receptors in the target alter the efficiency, although the wait time distribution remains unchanged in both cases. This provides constraints on the fusion mechanism and the identity of the rate-limiting step. Together, our data paints a picture of Sendai virus as a comparatively inefficient and slow fusion “machine” and sets the stage for investigation of other paramyxoviruses.

## Introduction/Background

Paramyxoviruses are a family of non-segmented negative-sense RNA, membrane-enveloped viruses that include many prominent pathogens, including human parainfluenza viruses, mumps virus, measles virus, and emerging pathogens such as Hendra and Nipah viruses (1). Sendai virus (SeV, formally murine respirovirus) is a prototypical member of the respirovirus genus of the *Paramyxoviridae* family (2, 3). While it causes animal disease and can even infect human cells, SeV does not cause human disease; therefore, it has been a unique candidate for various biotechnological applications.

As with other membrane-enveloped viruses, the initial stages of SeV infection are first, binding to a receptor on the host cell membrane, and second, fusion of the viral and host membranes to deliver the viral RNA (1, 4, 5). Both processes are mediated by viral proteins. The viral HN attachment protein binds to the receptor, an α2,3-linked sialic acid glycoprotein or glycolipid. This binding initiates a conformational change in the HN protein which then triggers the viral F protein to catalyze membrane fusion.

Although the key players in these processes have been identified, many central questions remain about the biophysical mechanism of fusion, such as the characteristic timescale of fusion, the numbers of proteins involved, the number and identity of rate-limiting steps in membrane fusion, etc. This is due in part to the fact that much of what is known about fusion comes from bulk or cell-based fusion studies (6–9), which typically convolve binding and fusion, and only allow observations of bulk viral behaviors.

Recently, researchers have begun using fluorescence microscopy measurements of single virions to study viral fusion, often using model lipid membranes as fusion targets (10–17). Such measurements enable precise control of the viral environment as well as deconvolution of binding and fusion processes. Coupled with mathematical modeling, these data provide a window into the timescale and number of rate-limiting steps and therefore into the biophysical fusion mechanism. Such approaches have been applied successfully to a variety of viral families (11, 12, 14, 17), although typically for viruses whose fusion trigger is environmentally manipulable (such as pH drop), and where binding and fusion are mediated by the same viral protein (such as hemagglutinin for influenza). There has not, to our knowledge, been any reported single virion fusion measurements of any paramyxovirus.

Here, we develop a single virus lipid mixing assay to study the fusion (lipid mixing) of SeV. We then apply that assay to interrogate fundamental questions about the biophysical mechanism of SeV fusion.

## Results/Discussion

### Single virus lipid mixing assay to study SeV

To our knowledge, there are no published reports of single virus fusion assays for any paramyxovirus. We therefore developed our single virus lipid mixing assay for SeV by adapting assays used for other viral families (11, 13–15) with some important modifications (full details in **Supplementary Material**). In our assay (see schematic in **Figure 1a**), the viral envelope is fluorescently labeled at a self-quenched level for visualization by fluorescence microscopy. The labeled virus is then injected into a microfluidic flow cell, in which liposomes have been tethered to a polymer surface via NeutrAvidin-biotin interaction. These liposomes are the fusion targets and contain gangliosides (either GQ1b or GD1a), which serve as the receptors for SeV. Binding of the labeled virions to these tethered liposomes occurs during a brief incubation (∼1 minute), following which unbound virions are rinsed from the microfluidic cell to reduce background noise. Bound virions are then observed by time-lapse microscopy. Additional characterization/validation measurements are discussed in the **Supplementary Material**, including validation that biotin-lipids did not alter SeV binding and characterization of SeV particles by immunofluorescence (**Figures S1-S2)**.

**Figure 1.**
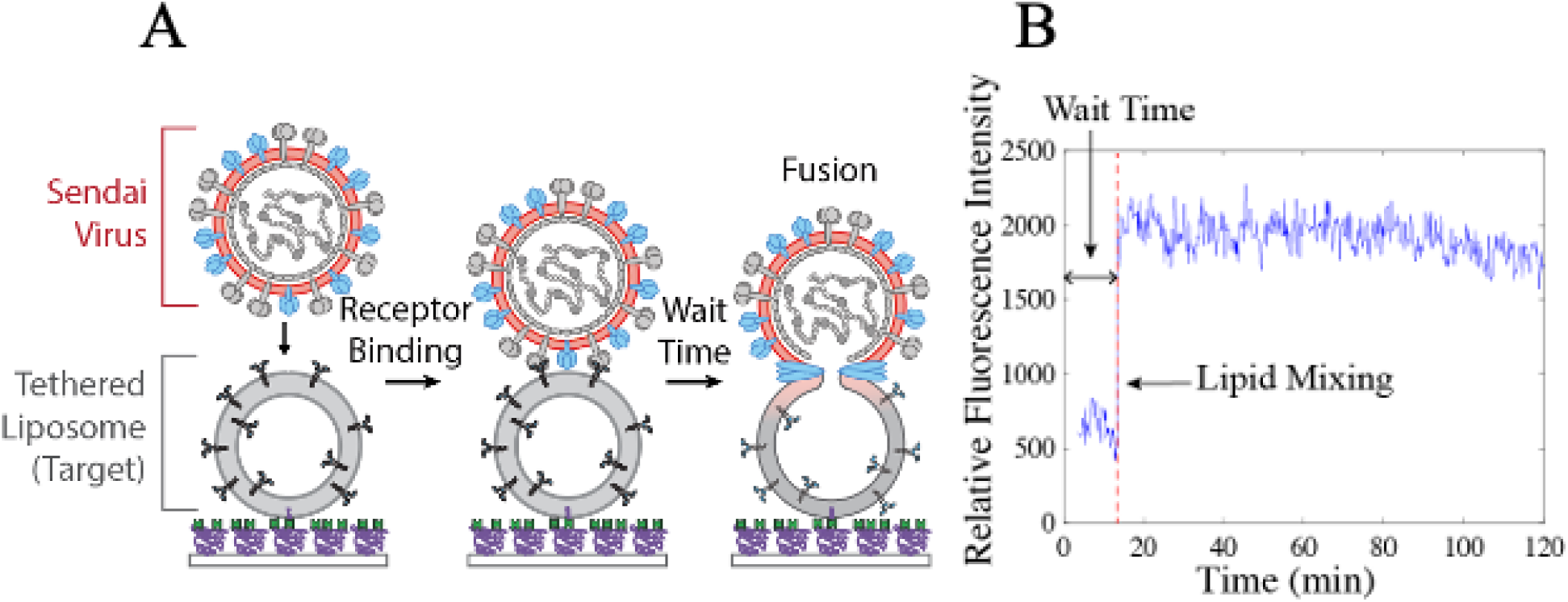
Overview of SeV single virus lipid mixing assay. A) Schematic of single virus assay. Target liposomes containing α2,3-linked sialic acid receptors are tethered to a PLL-PEG coated coverslip in a microfluidic flow cell via Neutravidin-biotin interactions. Fluorescently labeled SeV particles are injected into the flow cell (t = 0) and bind to the receptors during a 1 min incubation. Unbound viruses are removed by rinsing. Bound viruses are monitored by fluorescence microscopy, and fusion is observed as fluorescence de-quenching of the lipophilic dye upon lipid mixing between the viral and target membranes. B) Example fluorescence intensity trace of lipid mixing between a virion and target liposome. Fluorescence intensity is measured as the fluorescence within a region of interest around the bound virion in each video frame. Lipid mixing is detected as a jump in the intensity due to de-quenching. The wait time is defined as the time between viral binding (set at t=0) and lipid mixing.

One important modification in our assay was the length of time over which the virions were observed. In most prior single virus fusion reports, nearly all fusion activity occurred in the first few minutes following triggering (10–16). However, very few SeV particles (∼1-2%) were observed to undergo lipid mixing during the first few minutes following binding (leftmost data in **Figure S4**). Instead, virions continued to undergo fusion for at least an order of magnitude longer. This necessitated longer data collection times (typically ∼90-120 min of time-lapse microscopy) as well as rigorous removal of unbound virions prior to data collection to reduce background noise to an acceptable level for automated analysis.

We observed a variety of viral behaviors in the time-lapse videos. Some virions underwent lipid mixing, detected by fluorescence de-quenching of the dye-labeled lipid in the viral envelope as it was diluted into the target liposome (**Figure 1b**). Some virions underwent no change during the time-lapse video (**Figure S5**); others were observed to detach from the target liposome (**Figure S5**). Such unbinding events may be due to stochastic unbinding of the receptor, neuraminidase activity of the HN protein, or both (18, 19). While unbinding events are of potential interest for future study, the focus of this report is on viral fusion. Therefore, virions that unbound were excluded from analysis, as they could not be observed during the entire video (typically ∼4-8% of total virions in our data).

Two key metrics are obtained from analysis of our fusion assay data: 1) the wait time between binding and lipid mixing for viruses that undergo fusion and 2) the extent of lipid mixing, defined as the fraction of virions observed to undergo lipid mixing during the observation window.

### Distribution of wait times reveals insights into SeV fusion mechanism

The distribution of wait times from many viruses can be visualized as a cumulative distribution function (CDF, such as **Figure 2**), which contains information about the number and timescale of rate limiting steps, and therefore the mechanism of fusion. The extent alone provides information on the fraction of viruses that can achieve fusion under the experimental conditions being examined. In conjunction with the CDF, the extent provides constraints on the mechanism of fusion, as has been demonstrated for other viral families (13, 14, 20).

**Figure 2.**
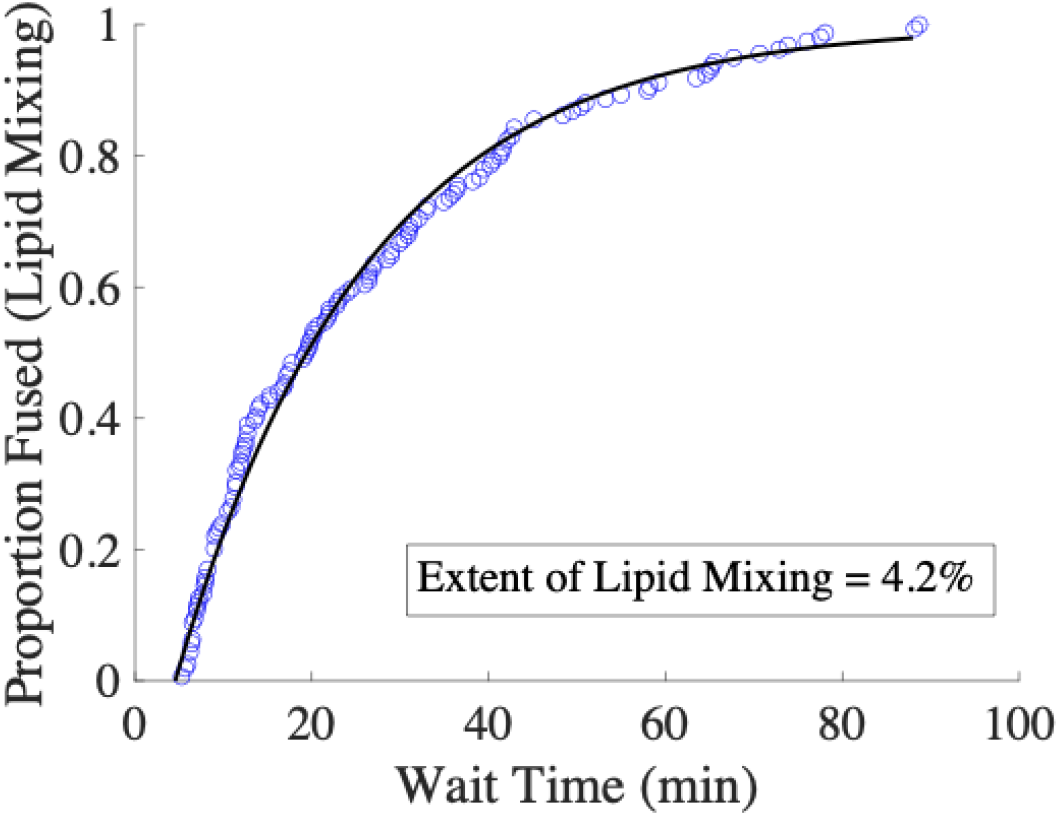
Cumulative distribution function (CDF) of SeV lipid mixing wait times follows a single exponential model. The distribution of wait times compiled from single virus lipid mixing measurements is shown as a CDF (blue circles), and modeled by a single exponential function with time lag (black line). The extent of lipid mixing was calculated as the fraction of virions that underwent lipid mixing during 90 minutes of observation (N = 159/3786). Lipid composition of target liposomes = 2% GQ1b, 20% DOPE, 30% cholesterol, 1% 16:0 biotinyl PE, 0.05% OG-DHPE, and 46.95% POPC.

Several features about the CDF and extent were notable in our SeV data. First, the observed distribution was exponential (**Figure 2**), fitting well to a single exponential curve with a time lag. This suggests that there is a single rate limiting step in the lipid mixing mechanism (*k* = 0.047 min^-1^). Second, as mentioned previously, the average timescale of lipid mixing was relatively long. For most viruses that have been studied using single virus measurements (typically being pH-triggered), average timescales of fusion are generally in the tens of seconds range (11–16). However, the average wait time for SeV lipid mixing was ∼20 minutes, and fusion continued even after 90 min of observation. Third, the extent of lipid mixing was rather low. Only ∼4% of viral particles were observed to undergo lipid mixing over 90 minutes. For comparison, reported maximum extents of fusion to model membranes for other viruses have been reported as: influenza virus = ∼30-80% (11, 15), Zika virus = ∼30% (14), West Nile virus = ∼30% (16), HIV = ∼50-60% (17).

Separately, we note that the initial segment of the CDF (wait time < ∼5 minutes) is not observed. This “dead time” represents the time between virus addition to the flow cell (t = 0) and the initiation of the time-lapse video (**Figures S3-S4b**), during which viral binding, rinsing of unbound viruses, and sample re-focusing occurs. This is necessary to reduce the background noise from unbound virions to manageable levels. In control measurements (**Figure S4**), we determined that the extent of fusion that occurs during our typical dead time was ∼1-2%. This matched well with the extrapolation of our single exponential fit to wait time = 0, indicating that the dead time is not obscuring a different curve shape or larger than expected change in extent.

### Trypsin treatment increases lipid mixing extent but doesn’t alter distribution of wait times

One explanation for the low extent of lipid mixing in our data (**Figure 2**) is a low fraction of properly processed F proteins. The SeV F protein is initially produced as a single polypeptide chain (the F_0_ form), which arranges into a homo-trimer. Following production, F_0_ must be proteolytically cleaved prior to subsequent infection; otherwise it cannot undergo the necessary structural rearrangements to catalyze fusion (1, 4). This cleavage divides each F protein into 2 fragments (F_1_ and F_2_), which remain covalently attached in the homo-trimer via disulfide bonds. Similar requirements for proteolytic processing are also observed for other viral families with class I fusion proteins (21).

Therefore, a low extent might be observed if the fraction of properly cleaved F proteins is also low, as many viruses might not have sufficient numbers of cleaved F proteins at the virus-target membrane interface to successfully achieve fusion. Additionally, depending on the identity of the rate-limiting step in the lipid mixing mechanism, this might also result in slower kinetics, leading to the long timescale of lipid mixing that we observed.

To investigate this, we asked whether trypsin treatment of SeV would alter the extent and/or kinetics of lipid mixing in our assay. Trypsin treatment of viruses (including SeV) has commonly been employed as a strategy to improve infection yields in cell culture, due to proteolytic cleavage of the fusion protein (22, 23).

In early tests, injecting TPCK-trypsin into the flow cell following virus binding did indeed appear to activate additional virions for lipid mixing. (**Figure S6**). Therefore, we conducted a set of single virus lipid mixing measurements in which we pre-treated SeV with different concentrations of TPCK-trypsin prior to injection into the flow cell (**Figure 3**). There were two notable observations. First, the lipid mixing extent was clearly sensitive to treatment with TPCK-trypsin. We observed a doubling in the extent from 10 µg/mL to 100 µg/mL TPCK-trypsin, after which a plateau appeared to be reached (**Figure 3b**). This demonstrates that trypsin treatment is sufficient to increase the extent, suggesting that the low initial extent was due, at least in part, to incomplete F protein processing. Second, the CDF kinetics were not substantially different between 0-100 µg/mL TPCK-trypsin – all curves were exponential in shape and of similar timescale. 1000 µg/mL TPCK-trypsin did exhibit a somewhat shifted CDF, possibly suggestive of proteolytic over-processing. Together, these results indicate that trypsin treatment up to 100 µg/mL does not alter the identity or timescale of the rate-limiting step.

**Figure 3.**
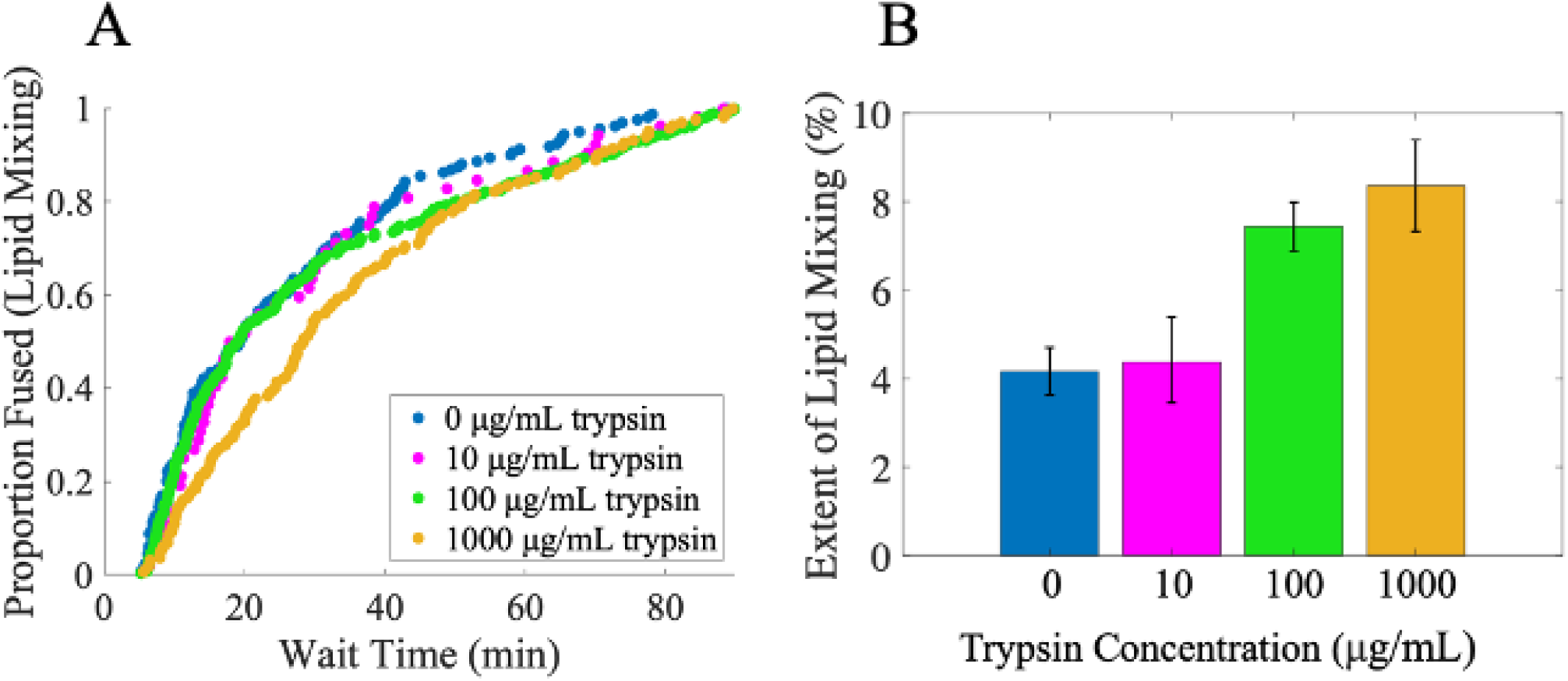
Trypsin treatment increases lipid mixing extent, with little change to CDF. Single virus lipid mixing experiments were conducted with SeV that had been pre-treated with various concentrations of TPCK-trypsin (0-1000 µg/mL). (A) shows the distributions of observed wait times. (B) shows the extents of lipid mixing, calculated as the fraction of observed virions that underwent lipid mixing during 90 minutes of observation. N_virus_: 0 µg/mL = 159/3786, 10 µg/mL= 52/1182, 100 µg/mL = 443/5907, 1000 µg/mL = 162/1929. Values shown are the mean ± 95% confidence interval determined by bootstrap resampling (NumBootstraps = 10,000). Extents at 100 µg/mL and 1000 µg/mL were statistically different from 0 µg/mL and 10 µg/mL with p < 0.0002 determined by bootstrap. All other extent comparisons were not significant. Lipid composition of target liposomes = 2% GQ1b, 20% DOPE, 30% cholesterol, 1% 16:0 biotinyl PE, 0.05% OG-DHPE, and 46.95% POPC.

In control experiments, we determined that trypsin treatment did not substantially alter lipid mixing extent during the experimental dead time (**Figure S4**). Additionally, we also determined that trypsin treatment did not substantially alter viral binding activity, indicating that the HN protein was unaffected by trypsin treatment (**Figure S7**).

Together, these results suggest that the density of properly cleaved F proteins at the virus-target interface is an important factor in determining the extent of lipid mixing but not the rate-limiting step. In turn, this suggests that the rate-limiting step is likely not dependent on the concerted action of multiple F proteins. Such a mechanism has been proposed for influenza virus (24), in which a minimum number of triggered fusion proteins at the virus-target interface is required for fusion, and adding additional activated fusion proteins further lowers the energetic barrier to fusion. If such a mechanism were employed by SeV, an increase in properly cleaved (and therefore activatable) F proteins should shift the CDF to shorter timescales. This however was not observed (**Figure 3**).

Separately, these results are of practical importance. Given that trypsin treatment increases the lipid mixing extent but leaves the rate-limiting step unaltered, we can use trypsin treatment to improve the throughput of our assay, allowing us to observe ∼2x the lipid mixing events per sample — especially important given the low baseline extent of the untreated virus.

### Influence of ganglioside receptor identity on SeV lipid mixing

Various α2,3-linked sialic acid glycoproteins and glycolipids can function as receptors for SeV *in vitro* (4, 6, 7). The effect of these receptors on viral binding has been well established, but their mechanistic influence on fusion is less well understood. It has been demonstrated that fusion and infectivity can be modulated by receptor identity (6–8, 25). However, prior work in this area has been done in bulk assays or cell culture, making it difficult to disentangle the effect on binding from fusion, particularly given that receptor binding is the trigger for fusion.

We previously investigated binding of SeV at the single-particle level to different ganglioside receptors, demonstrating that there is markedly different binding activity to different gangliosides (19). Of particular note were the gangliosides GD1a and GQ1b (chemical structures in **Figure S8**), with GQ1b yielding twofold higher binding but a less-steep cooperative binding response, indicative that fewer HN-GQ1b complexes may be required for stable binding relative to HN-GD1a complexes. Our data and other prior work (18) also suggest that individual HN-receptor complexes are highly dynamic; stable virus binding is determined by avidity rather than monomeric HN-receptor affinity.

To assess the effect of receptor identity on SeV fusion separately from its effect on binding, we conducted our single virus lipid mixing assay, observing fusion to target vesicles containing either GD1a or GQ1b (**Figure 4**). Interestingly, we observed that the extent of lipid mixing increased marginally with GQ1b relative to GD1a, but there was no significant difference between the CDFs. This suggests that receptor identity does not alter the rate-limiting step (else the CDF would shift), but does influence the fraction of viruses that ultimately get triggered for fusion. In turn, this suggests that GQ1b is more effective than GD1a at initiating F protein activation by HN, yielding a higher density of activated F proteins which then results in a higher extent. This is especially true if more HN-GD1a complexes are required for the virus to remain stably bound. However, this receptor-mediated activation is unlikely to be the rate-limiting step, else we might expect to see a faster shift in the CDF for GQ1b, which was not observed.

**Figure 4.**
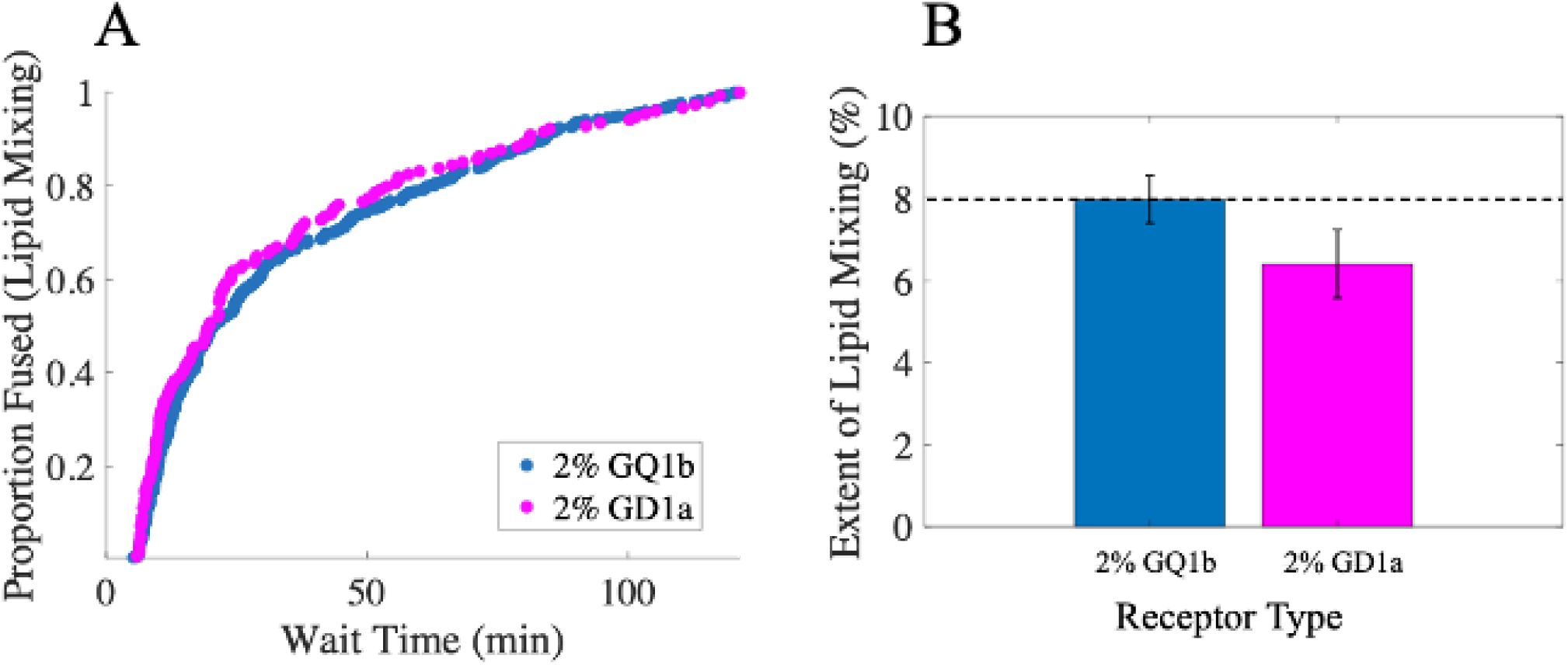
Influence of receptor identity on SeV lipid mixing. Single virus lipid mixing experiments were conducted to target liposomes with different receptors: either 2% GQ1b or 2% GD1a. SeV had been pre-treated 100 µg/mL TPCK-trypsin. (A) shows the distributions of observed wait times. (B) shows the extent of lipid mixing for each receptor, calculated as the fraction of observed virions that underwent lipid mixing during the experiment. N_virus_ for extents were: 2% GQ1b = 474/5925, 2% GD1a = 154/2406. Values shown are the mean ± 95% confidence interval determined by bootstrap resampling (NumBootstraps = 10,000). Extents between the 2 conditions were statistically different with p <.02 determined by bootstrap. Lipid composition of target liposomes = 2% GQ1b or GD1a, 20% DOPE, 30% cholesterol, 1% 16:0 biotinyl PE, 0.05% OG-DHPE, and 46.95% POPC.

## Conclusion

We have developed a single virus fusion (lipid mixing) assay for Sendai virus, the first reported single virus fusion measurement of any paramyxovirus to our knowledge. Using our assay, we found that the distribution of wait times was exponentially distributed, suggesting a single rate limiting step in the lipid mixing mechanism. Surprisingly, the distribution was also very slow, with average fusion wait times of ∼20 min after viral binding. The extent of lipid mixing (fraction of viruses observed to undergo fusion) was also rather low: ∼4% for untreated virus over 90 min. Together, this paints a picture of SeV as a somewhat inefficient fusion “machine” compared to other viral families, with typical average wait times ∼tens of seconds and extents of ∼30-80% (11–17).

Treatment of SeV with trypsin caused a doubling in the extent, suggesting that the low extent was due in part to incomplete cleavage of F fusion proteins on the viral surface. However, trypsin treatment produced little change to the distribution of wait times, suggesting that the rate-limiting step does not depend on the concerted action of multiple fusion proteins, as has been suggested for influenza virus (24).

We also examined the influence of the target receptor on the lipid mixing kinetics, and found that while GQ1b marginally increased the extent of lipid mixing compared to GD1a, the distribution of wait times remained unchanged. Together, this suggests that the density of activated F proteins can influence the extent of lipid mixing, but that receptor-mediated activation itself is unlikely to be the rate-limiting step.

It is unknown how many F protein trimers are necessary to effectively catalyze membrane fusion for SeV or other paramyxoviruses, although for other viral families the minimum number of required fusion proteins ranges from ∼2-5 (11, 12, 14, 20, 24), with models often requiring activated fusion proteins to be spatially adjacent. Our data does not rule out the possibility that multiple (possibly adjacent) F proteins may be required to achieve lipid mixing, serving as a gate-keeping mechanism which would control the extent. However, both our trypsin and receptor data (**Figures 3 and 4**) suggest the rate-limiting step itself does not appear to depend on the action of multiple fusion proteins, or on their activation.

Finally, we note that even after trypsin treatment, the extent of fusion remained rather low (<10%). This may suggest that very few virions are fusion-competent to begin with. Physiological data on infectious unit-to-particle ratios are sparse, and we are not aware of such data for any paramyxovirus. Such data does exist for dengue virus (26), with infectious unit-to-particle ratios in the range of 1:several thousand, with only 1 in 6 particles being fusion competent, indicating that similarly low viral fusion extents are possible for other viral families. It is even possible that fusion incompetent virions may serve a useful role for the virus in host pathogenesis or immune evasion, similar to the role played by partially mature dengue viruses (27), although this remains speculation for SeV.

## Acknowledgments

The authors gratefully acknowledge financial support from Williams College and NIH grant R15AI171754. The authors thank Grayson Brooks and Caroline Hon (Williams College) for useful discussions.

## Author Contributions

LJ, DY, AP, and KB designed experiments, collected and analyzed data. LJ also helped write the manuscript. RJR designed experiments, analyzed data, acquired project funding, provided supervision, and wrote the manuscript. All authors reviewed the manuscript.

