## Supplementary Material for "Single virus lipid mixing study of Sendai virus provides insight into fusion mechanism"

**Correspondence**

### S.0: Supplementary Materials and Methods

#### Materials

Dioleoylphosphatidylethanolamine (DOPE), palmitoyloleoylphosphatidylcholine (POPC), cholesterol (Chol), 1,2-dipalmitoyl-sn-glycero-3-phosphoethanolamine-N-biotinyl (biotinyl PE), and ganglioside receptors GD1a and GQ1b were purchased from Avanti Polar Lipids (Alabaster, AL, USA). Neutravidin protein, Oregon Green-1,2-dihexadecanoyl-sn-glycero-3-phosphoethanolamine (OG-DHPE) and Texas Red-1,2-dihexadecanoyl-sn-glycero-3-phosphoethanol-amine (TR-DHPE) were obtained from Thermo Fisher Scientific (Waltham, MA, USA). Poly-L-Lysine graft PEG polymer (PLL(20)-g[3.5]-PEG(2), abbreviated PLL-g-PEG) and Poly-L-lysine graft biotinylated PEG polymer (PLL(20)-g[3.5]- PEG(2)/PEG(3.4)-biotin(50%), abbreviated PLL-g-PEG-biotin) were obtained via SuSoS AG (Dübendorf, Switzerland). Polydimethylsiloxane (PDMS) Sylgard 184 elastomer base and curing agent were acquired from Ellsworth Adhesives (Germantown, WI, USA). Chloroform, methanol, and buffer salts were purchased from Fisher Scientific (Pittsburgh, PA) and Sigma-Aldrich (St. Louis, MO, USA). Bovine serum albumin (BSA) was purchased from Sigma Life Science (Darmstadt, Germany). TPCK-Trypsin was purchased from Thermo Fisher Scientific (Waltham, MA, USA). Sendai virus (purified Sendai Cantell Strain, egg-grown, batch 960216) was acquired from Charles River Laboratories (Wilmington, MA, USA) and was handled following a BSL-2 protocol at Williams College. Anti-Sendai Virus HN protein (1A6) mouse IgG2a antibody was acquired via Kerafast Inc (Boston, MA, USA) from the laboratory of Prof. Benhur Lee, Icahn School of Medicine at Mount Sinai, NY, USA. Goat anti-mouse IgG-Alexa 488 antibody was obtained from Abcam (Cambridge, UK).

#### Buffer Definitions

- Reaction buffer (RB pH 7.4) = 10 mM NaH2PO4, 90 mM sodium citrate, 150 mM NaCl, pH 7.4.
- HEPES Buffer (HB pH 7.2 or 8.0) = 20 mM HEPES, 150 mM NaCl, pH 7.2 or 8.0.

All buffers were titrated to the indicated pH with HCl or NaOH, then filtered at 0.22 µm pore size.

#### Fluorescence Microscopy

Fluorescence microscopy was performed using a Zeiss Axio Observer 7 microscope (Carl Zeiss Microscopy, LLC., White Plains, NY) with a 63x oil immersion objective, NA=1.4 , a Lumencor Spectra III, LED Light Engine, and a Definite Focus 3. A Hamamatsu ORCA Flash 4.0 V2 Digital CMOS camera (Hamamatsu Photonics K.K., Hamamatsu City, Japan) with a 16-bit image setting was used to capture images, in conjunction with Micromanager software (1). Fusion timelapse videos were acquired at 100 ms/frame and 2x2 binning, while continuous video micrographs were taken at 300 ms/frame with 2x2 binning. Filter cube settings for Texas Red images were: ex = 562/40 nm, bs = 593 nm, em = 641/75 nm, light engine typical intensity setting = 10/1000 or 25/1000 (green LED). The excitation/emission filter cube for Oregon Green and Alexa 488 images was: ex = 475/50 nm, bs = 506 nm, em = 540/50 nm, light engine typical intensity setting = 2/1000 (cyan LED).

**Labeling and Concentration Estimation of Sendai Virus**

Labeling and concentration estimation were performed as previously described (2). Briefly, to label Sendai virus, Texas Red-DHPE (0.75 g/L in ethanol) was mixed with HB pH 7.2 in a 1:30 ratio. The dye-buffer mixture was sonicated and heated to 60℃ for 30 minutes, then cooled before the addition of virus. 120 μL of the dye buffer mixture was added to each 30 μL aliquot of unlabeled virus, and the SeV mixture was incubated for 2 hours at room temperature to allow the lipophilic dye to incorporate into the viral membrane. 1300 μL of HB pH 7.2 was added to each aliquot following incubation, and the mixture was centrifuged at 21K x g at 4℃ for 50 min to isolate labeled viral particles from free dye in the solution. The supernatant containing the free dye was removed, and the pellet was resuspended in HB pH 7.2. The virus was then filtered through a Millex-GV 0.45 μm PVDF low binding filter (MilliporeSigma, Burlington, MA, USA) to remove dye and virus aggregates, then stored at -80℃ until use. Viral protein content was assessed via BCA assay and used to estimate viral particle concentration as we have previously described (2).

**Preparation and Activity Verification of TPCK-Trypsin**

Powdered TPCK-treated trypsin (stored at -20℃) was dissolved in HB pH 8.0 at 4000 μg/mL, then filtered using a Millex-GV 0.22 μm PVDF low binding filter. After separation into single-use aliquots, the dissolved trypsin was stored at -20℃ until use. Trypsin activity was verified using a routine colorimetric assay monitoring the hydrolysis of p-nitrophenyl acetate (3).

**Preparation of Microfluidic Flow Cell**

Microfluidic flow cells were constructed using polydimethylsiloxane (PDMS) pieces plasma-bonded to cleaned glass coverslips, as previously described (2). Briefly, PDMS pieces consisted of two parallel flow channels, each with dimensions 2.5 mm x 13 mm x 70 μm, an approximate volume of 4 μL, and 2.5 mm diameter inlet/outlet holes. PDMS was composed of a 10:1 mixture of Sylgard 184 elastomer base and curing agent, which was degassed under house vacuum for 1 hour before being poured into a mold, then cured at 70℃ for a minimum of 2 hours. Following the curing process, a scalpel was used to separate individual PDMS pieces from the mold, and inlet/outlet holes were punched into the ends of the channels with a biopsy hole puncher (2.5 mm diameter, Harris Uni-core, Ted Pella Inc.). To clean glass coverslips, (24 x 40 mm, NO. 1.5 VWR International, Randor, PA), a 1:7 solution of 7X detergent (MP Biomedicals, Burlingame, CA) and deionized water was prepared. The coverslips were fully submerged in the solution, then heated until the solution turned clear. After extensive rinsing under deionized water for a minimum of 2 hours and MilliQ water for 2 minutes, the coverslips were heated to 400°C in a kiln for 4 hours. Both PDMS chips and cleaned glass coverslips were stored at room temperature and used within approximately one month. To plasma-bond PDMS chips to coverslips, a Harrick Plasma Cleaner PDC-3xG (Harrick Plasma, Ithaca, NY) was used to activate the bonding surfaces of a PDMS chip and coverslip via exposure to air plasma for 70 seconds. Following assembly of the flow cell, buffer wells, made by cutting the ends off 1 mL plastic pipette tips, were glued over the inlet holes using five-minute epoxy (Devcon, ITW Polymer Adhesives North America, Danvers, MA).

**Preparation of Target Liposomes**

Target liposomes were prepared using a thin-film hydration and extrusion method, as previously described (2). Briefly, lipids in chloroform/methanol were added to a test tube with a glass syringe at the desired molar ratio (1.4 x $10^{-7}$ moles of total lipid) and dried down to a film using a stream of $N_2$ gas. Unless otherwise noted, the standard liposome composition was 2% ganglioside receptor, 20% DOPE, 30% cholesterol, 1% 16:0 biotinyl PE, 0.05% OG-DHPE, and 46.95% POPC. The lipid film was further dried under house vacuum for a minimum of two hours to remove trace solvent. 250 µL of RB pH 7.4 was added following desiccation to rehydrate the lipid mixture for approximately ten minutes, which was then vortexed at maximum speed for a minimum of sixty seconds to ensure complete recovery (4). The resuspended lipid mixture was passed 21 times through a mini-extruder (Avanti Polar Lipids, Alabaster, AL) with a 100 nm-pore membrane to generate large unilamellar liposomes. The resulting liposomes were stored at 4°C for use within approximately one week.

**Assembly of Target Liposome Surface in Microfluidic Flow Cell**

Biotin-NeutrAvidin binding between biotinylated lipids in liposomes and a biotinylated polymer was used to tether liposomes. After assembly of the flow cell as described above, 5 µL of a 95:5 PLL-PEG:PLL-PEG-biotin mixture was added to the inlet of each channel and incubated for 30 minutes at room temperature. A Fusion 200 syringe pump (Chemyx Inc., Stafford, TX, USA) with a flow rate of 800 µL/min was used to rinse each channel with 1 mL of MilliQ water followed by 1 mL of RB pH 7.4. Excess buffer was removed from the outlet and inlet holes before exchanging 6 µL of 0.2 mg/mL NeutrAvidin protein dissolved in RB pH 7.4 into the channels. The flow cell was subsequently incubated for 15 minutes before repeating the rinsing process with 2 mL of RB pH 7.4. After removing excess buffer, 6 µL of target liposomes with biotinyl PE (0.4 mg/mL total lipids) was exchanged into each channel and incubated for one hour. The tethered liposome surface was rinsed once again with 2 mL of RB pH 7.4 before imaging to verify the quality of the surface.

**Single Virus Fusion (Lipid Mixing) Assay**

After imaging the tethered liposome surface, excess buffer from the rinsing process was removed from the inlet/outlet and 5 µL of Sendai virus (typical concentration = ~20 µg/mL total viral protein or ~40 pM viral particle concentration) was injected into the channel. If treated with trypsin, Sendai virus and dissolved TPCK-trypsin (4000 µg/mL) were thawed on ice prior to treatment. The trypsin was diluted with HB pH 8.0 to the desired concentration, then mixed with SeV in a 1:1 ratio and incubated at 37℃ for 20 minutes before addition to the channel. After a minute-long incubation at 37°C to allow for viral binding, 1 mL of RB pH 7.4 pre-incubated at 37°C was used to rinse out unbound virions. The rinse was performed via syringe pump, flow rate 800 µL/min. The sample was refocused, the timelapse was resumed in the Texas Red channel. Timelapse data was collected for 90 to 120 minutes at 37°C, with 7.5 second intervals between frames unless otherwise noted.

**Data Analysis and Model Fitting**

Timelapses were analyzed using custom Matlab scripts (source code at https://github.com/rawlelab/Sendai-Fusion-Analysis-Public), adapted from those that have been previously described (5). The scripts locate bound virions, track fluorescence intensity over time, identify lipid mixing events via dye dequenching, which is indicated by a sudden jump in fluorescence intensity, and calculate the wait time between binding and lipid mixing. An important aspect of the wait time calculation is determining the time at which binding occurs. Given the very short binding window (~1 minute) relative to the length of the time-lapse video (~90-120 min), the binding time of all virions (t = 0) was set at the time of introduction of the virions into the flow cell. This enabled more straightforward automated analysis by finding and monitoring the bound virions after the excess virions had been removed from the flow cell. Ultimately this introduces an uncertainty of ~1 minute to our wait time calculations. Automatic classification of fusion events by the script was verified manually by the user and corrected if necessary.

Model fits to CDFs were performed in Matlab using maximum likelihood estimation. To account for experiment-to-experiment variance (typically 3.5-5 min) in the "dead time" between addition of virus and the onset of the timelapse, a 5 minute cutoff was applied to the beginning of each CDF.

The single exponential model used was defined as:

$$F(t) = 1 - e^{-k\left(t - t_{lag}\right)}, \; t \geq 0 \qquad \textbf{[Equation S1]}$$

where $F(t)$ is the cumulative probability of lipid mixing at wait time $t$, $k$ is the rate constant, and $t_{lag}$ is the time lag, included to account for the "dead time". $k$ can also be expressed as $1/\tau$, where $\tau$ is the half-life decay time.

**Measurement of Fusion Events During Typical "Dead Time"**

To determine the extent of lipid mixing during the typical "dead time" between addition of virus and initiation of the timelapse, the minute-long incubation and rinsing steps were eliminated from the single virus fusion assay procedure described above. Instead, immediately after addition of virus, ~0.5 mL of pre-incubated buffer was added to the buffer well and a 10 minute-long continuous video micrograph was initiated. Due to the high background noise of unbound virions, video micrographs were analyzed manually in FIJI. 100 virions were selected at random and monitored for lipid mixing (fluorescence dequenching) during 5 minutes, contingent on the virion binding and remaining visible in the video during the entire 5 minute window.

**Single Virus Binding Assay**

Single virus binding assays were performed to supported lipid bilayers as previously described (2). Briefly, a flow cell was constructed following the procedure described above, and a supported lipid bilayer was formed in each channel via the vesicle fusion method. After extensive rinsing to remove unbound vesicles, 5 µL of Texas Red-labeled SeV (pre-treated or not with TPCK-trypsin) was injected into the channel and incubated for 15 minutes at room temperature. Unbound virions were removed by syringe pump rinsing (1 mL RB at 800 µL/min), and images of bound virions were collected throughout the flow cell. The number of viral spots in each image was quantified using custom-built Matlab scripts that have been previously described (2), and available at https://github.com/rawlelab/SendaiBindingAnalysis.

**Immunofluorescence (IF) Measurements**

IF experiments were performed as previously described (2). Briefly, 4 µL of virus was added to a channel, either a naked glass surface for non-specific binding, or a tethered liposome surface for specific binding. Virions were bound during a 15 min room incubation at room temperature, followed by surface passivation with 4 µL of 30 mg/mL BSA in HB pH 7.2 for 15 minutes. Then, 4 µL of 1:50 dilution of 1A6 Anti-HN primary antibody in HB pH 7.2 was added and the flow cell was incubated for 2 hours. The flow cell was then rinsed with 2 mL of HB pH 7.2, and passivated with 4 µL of BSA for 15 min. Goat Anti-Mouse IgG (AlexaFluor 488) secondary antibody (1:1000 dilution in HB pH 7.2) was added and incubated for 15 min. The channel was rinsed with 3 mL HB pH 7.2, then a given location in the channel was sequentially imaged in the Alexa 488 and Texas Red channels. The resulting images were analyzed and the fraction of bound virions that were co-localized with antibody (Alexa 488) signal was determined via custom-built Matlab scripts that have been previously described (2), and available at https://github.com/rawlelab/SendaiBindingAnalysis.

### S.1: Single SeV Lipid Mixing Assay Validation/Characterization Measurements

Single virus assays have been well-validated for other viral families, but they have not been utilized for any paramyxovirus (to our knowledge), and they have most often been used to observe fusion of viruses whose fusion trigger is pH or another environmental condition, not viruses whose fusion trigger is receptor binding. Therefore, we performed additional measurements to validate/characterize this assay for SeV.

First, we investigated the effect of including biotin-labeled lipids in the target vesicles on the viral binding process. We and others have previously demonstrated that SeV binds specifically to GD1a and GQ1b (2, 6, 7), the ganglioside receptors used in this report. To verify that the addition of biotin did not increase non-specific binding, we examined virus binding to supported lipid bilayers with and without biotin-labeled lipids and GQ1b, using a single virus binding assay we have reported previously (see Ref (2) and brief description in Supplementary Materials and Methods). We observed that the inclusion of biotin-labeled lipids in the target membrane did not induce non-specific virus binding to any degree (see **Figure S1**).

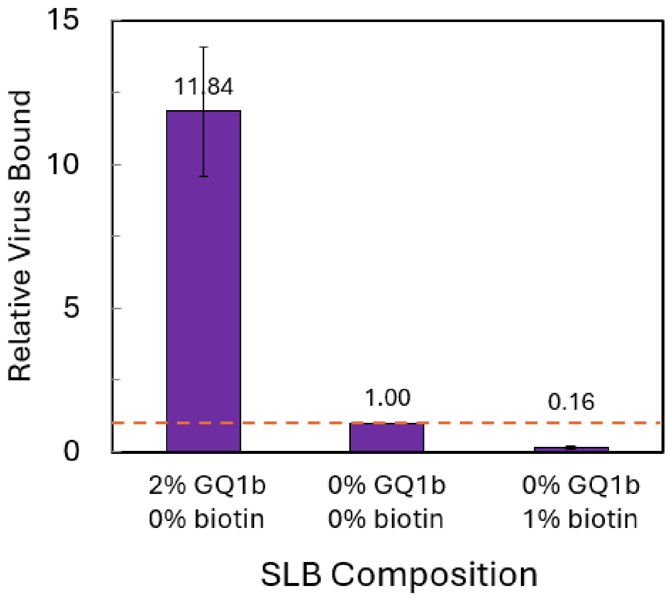

***Figure S1. Inclusion of biotin-lipid in target membranes does not induce non-specific binding of SeV.*** *Single-virus binding measurements were performed to supported lipid bilayers that contained the listed combination of biotin-DPPE and/or the viral receptor GQ1b. Binding is calculated relative to the no receptor, no biotin condition. Error bars are ± standard deviations of 5+ separate image locations within each sample, with a propagated relative error as described previously (2). Lipid composition of target SLBs = 0 or 2% GQ1b (as listed), 0 or 1% biotin-DPPE (as listed), 20% DOPE, 10% cholesterol, 0.05% Oregon Green-DHPE, and the remainder POPC (67.95%–69.95%).*

Second, we characterized the fraction of putative viral particles that bind to tethered vesicles which are indeed Sendai virions. Previous work has suggested that other cellular particles may be co-purified with viral particles even under rigorous purification schemes (8). We utilized an immunofluorescence (IF) assay to characterize the putative viral spots we observe of fluorescently-labeled SeV particles which are bound specifically to target vesicles containing 2% GQ1b receptor, or nonspecifically to either a clean glass surface. The antibody we utilized targeted the HN attachment protein (1A6 antibody, which has been reported previously, (2)). We observed that the fraction of viral particles that were IF-positive was very similar between particles bound to the target vesicles with GQ1b receptor, or nonspecifically to a clean glass surface (fraction IF-positive = ~0.5, see **Figure S2**). This suggests that the viral particles bound to the target vesicles containing GQ1b receptor are representative of the labeled viral particles as a whole. We note that the fraction IF-positive is a minimum bound on the fraction of particles that are indeed Sendai virions – we cannot rule out that the particles which did not test positive by IF could simply be virions that are insensitive to detection by the assay.

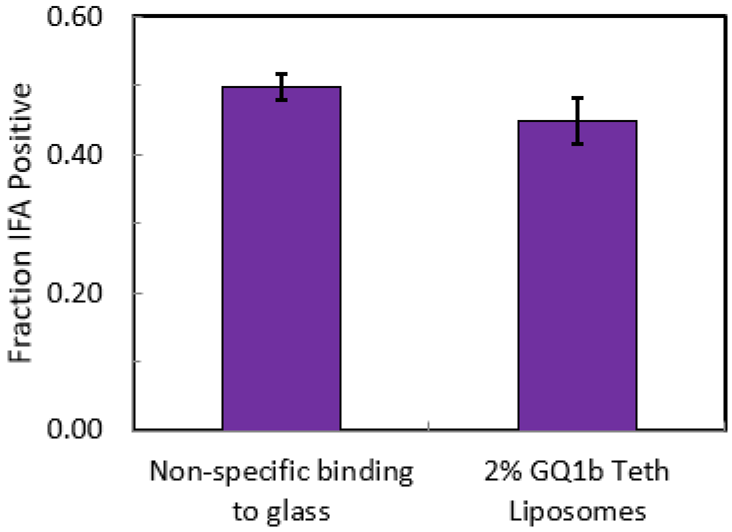

***Figure S2. Immunofluorescence measurements of SeV bound to tethered vesicles compared to non-specifically bound SeV.*** *SeV labeled with TR-DHPE was bound either non-specifically to a clean glass surface, or specifically to target liposomes containing 2% GQ1b. Bound SeV particles were sequentially treated with 1A6 anti-HN primary antibody and Alexa 488-labeled secondary antibody. Fraction IFA positive was calculated as the average number of TR-labeled particles in fluorescence micrographs that were colocalized with Alexa 488 signal. Values shown are average ± standard deviation of fraction IFA positive measured from 10 separate locations for each condition. Tethered liposomes contained 2% GQ1b, 20% DOPE, 30% cholesterol, 1% 16:0 biotinyl PE, 0.05% OG-DHPE, and 46.95% POPC.*

### S.2: Measurement of Fusion Events During Typical "Dead Time"

As described in the main text, the initial segment (wait time < several minutes) of our CDFs is not observed. This "dead time" represents the time between addition of the virus to the flow cell (t = 0) and the initiation of the time-lapse video (example trace with labeled dead time shown in **Figure S3**). During this time, the virions are allowed to bind for 1 min, following which unbound viruses are rinsed from the flow cell, and the sample is re-focused. This entire process takes several minutes, and is necessary to properly reduce the background noise from unbound viruses to manageable levels for automated analysis of the subsequent time-lapse video, which is collected for 90-120 min.

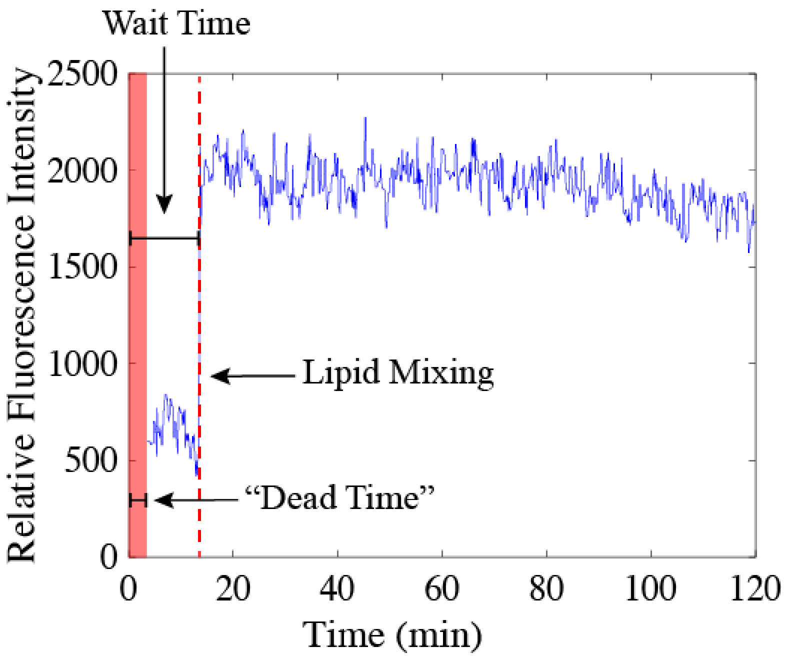

***Figure S3. Example fluorescence intensity trace of lipid mixing between a SeV virion and target liposome with dead time indicated.*** *The fluorescence intensity is measured as the fluorescence within a region of interest around the bound virion in each video frame. Lipid mixing is detected as an abrupt jump in the intensity due to fluorescence de-quenching. The wait time is defined as the time between viral binding (set at t=0) and the onset of lipid mixing. The dead time is the time between addition of the virus to the flow cell and the initiation of the time-lapse video.*

It is possible to collect data with zero dead time if the sample is not rinsed following virus injection and the sample is monitored via continuous (as opposed to time-lapse) video microscopy (see Measurement of Fusion Events During Typical "Dead Time" section in the Materials and Methods). In this case, viruses which are already bound at the beginning of the video are ignored, but viruses which are observed to bind during the video are monitored for lipid mixing. However, the throughput (number of viruses observed per sample) of these experiments is rather limited and the observation window must be limited to several minutes. Otherwise, the background noise from the excess unbound or loosely bound virions becomes overwhelming. Even still, the background noise is high enough that manual rather than automated analysis must be performed.

To quantify the extent of fusion that occurs during our typical dead time, we collected data using this "zero dead time" approach, and measured the extent of fusion over 5 minutes following virus binding. Data was collected for SeV untreated or treated with 100 µg/mL TPCK-trypsin. For both conditions, we observed an extent of ~2% during the first 5 minutes (blue bars in **Figure S4A**). We then compared this data to model predictions generated from the corresponding CDF data (i.e. **Figure 2** and green CDF in **Figure 3** in the main text). The model predictions were generated by extrapolating the single exponential fit from each CDF data back to wait time = 0 to calculate the "missing" proportion fused during the dead time and then multiplying by the observed extent of lipid mixing at 90 min (see example in **Figure S4B**). We found that the model predictions fell within the measurement error of the "zero dead time" data ( **Figure S4A**).

Taken together, this further supports the application of the single exponential model to the CDF data, and indicates that the dead time is not obscuring a larger than expected change in the extent, validating the data collection approach used in the remainder of this report.

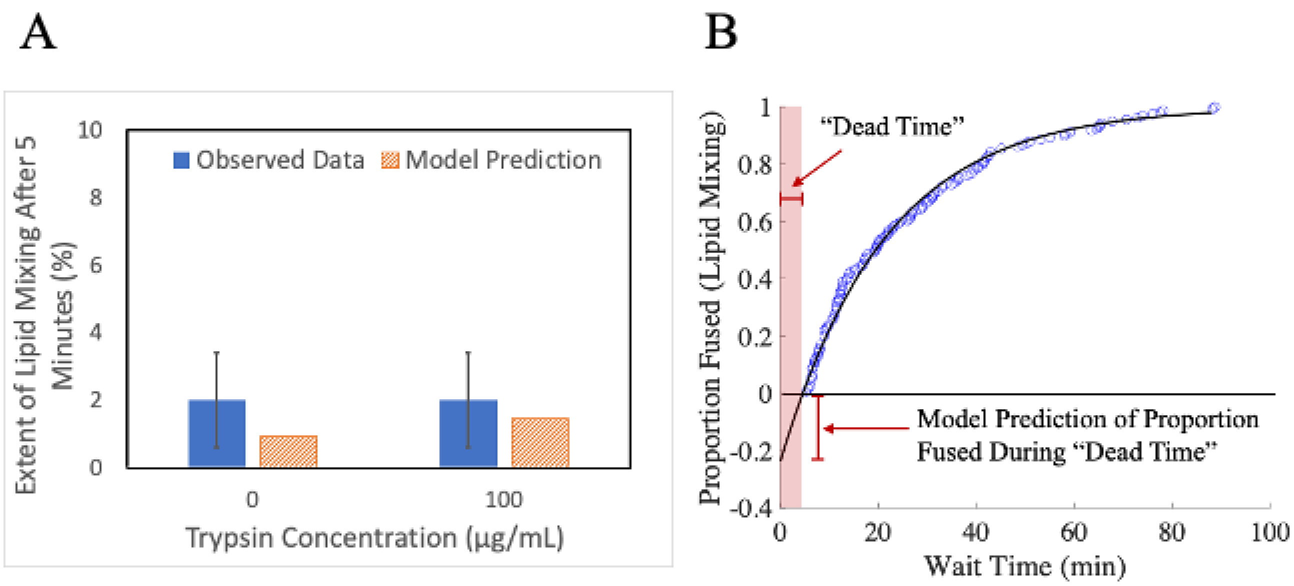

***Figure S4. "Fast" timescale measurements of lipid mixing extent match predictions from exponential fit model to CDFs.*** *A) Single virus lipid mixing experiments were conducted with untreated SeV or SeV that had been pre-treated with 100 µg/mL TPCK-trypsin. However, the flow cell was not rinsed following virus addition, and data collection was initiated immediately using continuous video microscopy (as opposed to time-lapse imaging). The extent of lipid mixing (observed data) was calculated as the fraction of observed virions that underwent lipid*

*mixing during the first 5 minutes following binding (Num Virus Observed = 100). Error bars are ± standard deviation determined by bootstrap resampling of the observed data (NumBootstraps = 10,000). Model predictions were determined from the CDFs of the longer timescale lipid mixing data (e.g. Figure 2 and Figure 3A in the main text). They were calculated by extrapolating the single exponential fit to each CDF back to wait time = 0 to determine the missing proportion fused. This was then multiplied by the observed extent of lipid mixing at 90 minutes (e.g. Figure 3B in the main text). B) Example extrapolation of single exponential model to wait time = 0 with missing proportion fused indicated. The CDF shown is the same as Figure 2 in the main text.*

### S.3: Example traces of different viral behaviors

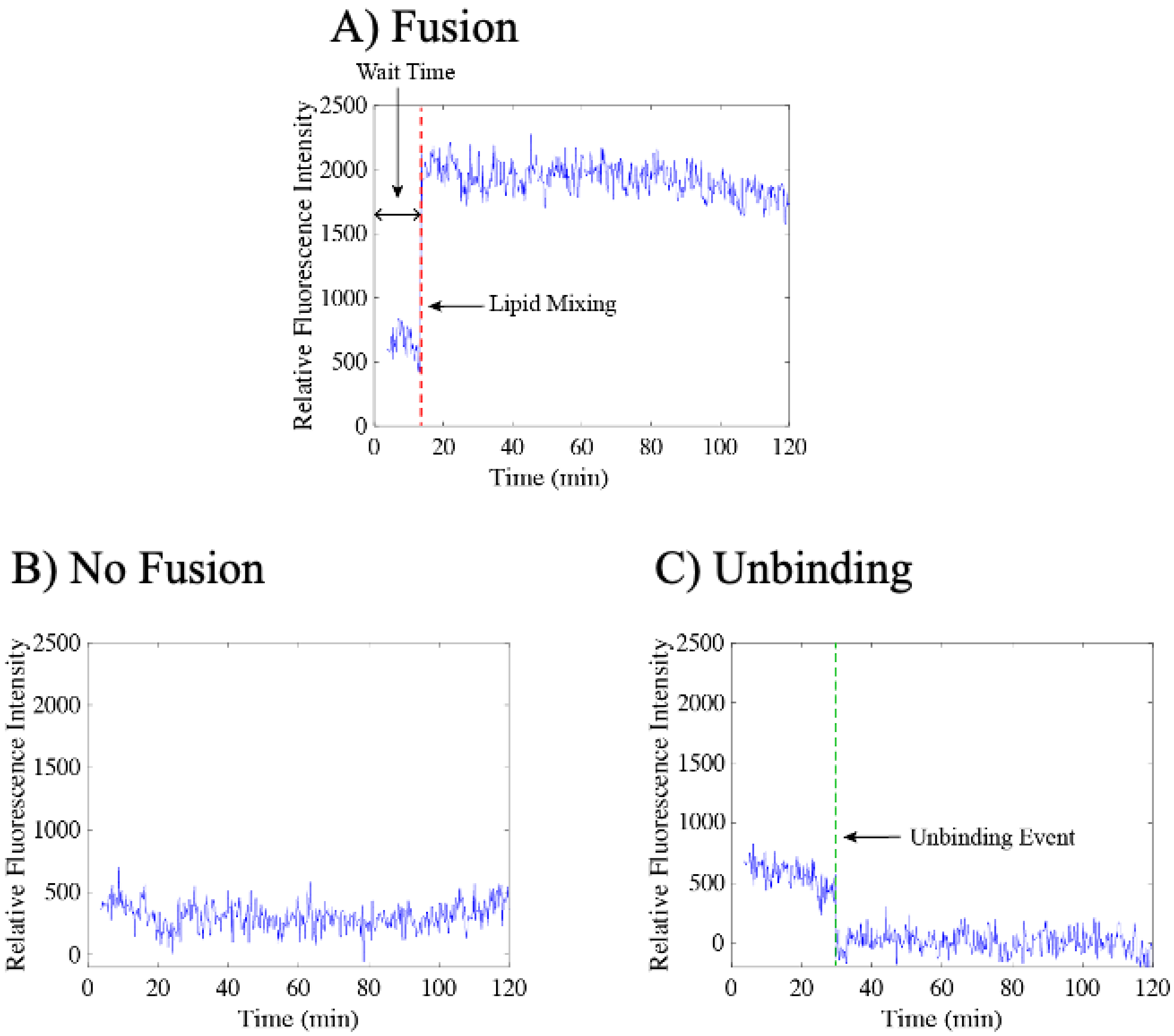

***Figure S5*** *Example fluorescence intensity traces of different observed behaviors of individual SeV virions bound to target liposomes. The fluorescence intensity is measured as the fluorescence within a region of interest around the bound virion in each video frame. (A) shows an example fusion (lipid mixing) event. Lipid mixing is detected as an abrupt jump in the intensity due to fluorescence de-quenching. The wait time is defined as the time between viral binding (set at t=0) and the onset of lipid mixing. (B) shows an example no fusion event, where the fluorescence intensity of the virion remains unchanged throughout the video. (C) shows an example unbinding event. Unbinding is detected as an abrupt decrease in the fluorescence intensity, returning to background intensity.*

### S.4: Trypsin treatment during viral lipid mixing measurement

As an initial test of the effect of trypsin treatment on viral fusion activity, we prepared a SeV single virus lipid mixing assay, and began collecting a time-lapse video. At the 10 minute mark, we injected 1000 µg/mL TPCK-trypsin into the microfluidic flow cell, and then continued the time-lapse video. We observed a marked increase in the rate of lipid mixing following trypsin addition (**Figure S6**). This was a clear proof of concept that proteolytic processing could alter the observed fusion behavior of SeV. However, this approach could not be used to quantitatively assess changes to either the extent or kinetics of lipid mixing, as the timescale of lipid mixing is convolved with the timescale of proteolytic processing.

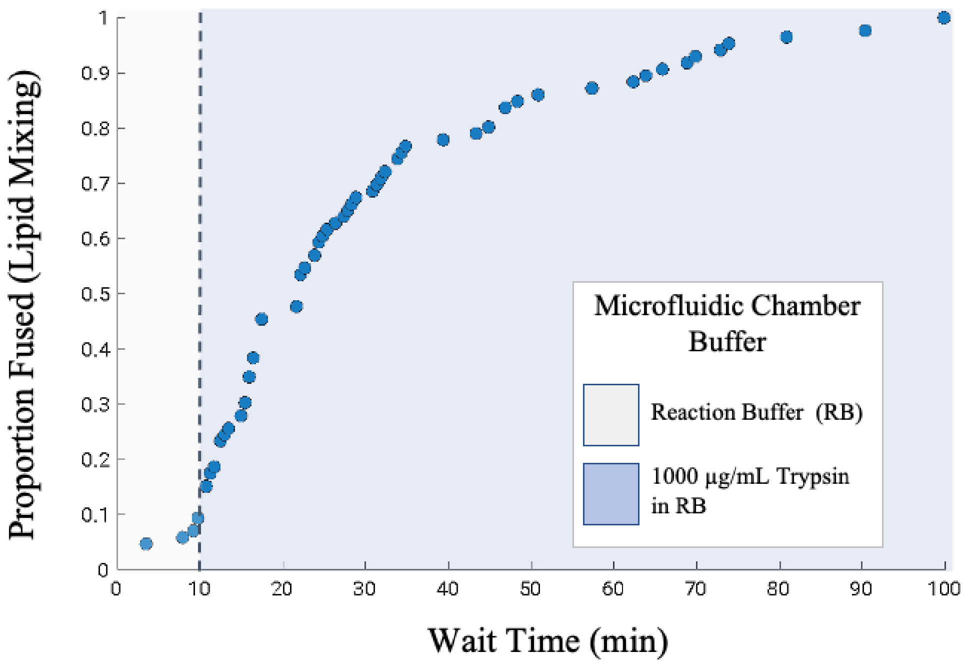

***Figure S6. Increase in lipid mixing is observed following injection of TPCK-trypsin.*** *A single virus lipid mixing experiment was initiated. At t = 10 min, 1000 µg/mL TPCK-trypsin was exchanged into the microfluidic chamber and the timelapse data collection was resumed.* $N_{virus}$ *for final extent of lipid mixing was 86/677. Lipid composition of target liposomes = 2% GQ1b, 20% DOPE, 30% cholesterol, 1% 16:0 biotinyl PE, 0.05% OG-DHPE, and 46.95% POPC.*

### S.5: Trypsin treatment doesn't alter viral binding activity

To assess whether our trypsin treatment may have inadvertently caused unintended alterations in other essential viral proteins, we conducted experiments to determine whether trypsin treatment had altered viral binding activity, which is governed by the HN protein, not the F protein. Using fluorescence microscopy, we quantified viral binding to supported lipid bilayers containing 2% GQ1b using a single virus binding assay we have reported previously (see (2) and brief description in Supplementary Materials and Methods). We compared the binding of untreated SeV to SeV treated with 100 µg/mL of TPCK-trypsin (**Figure S7**). We observed no substantial difference in binding between the two conditions, indicating that trypsin treatment had not altered viral binding activity

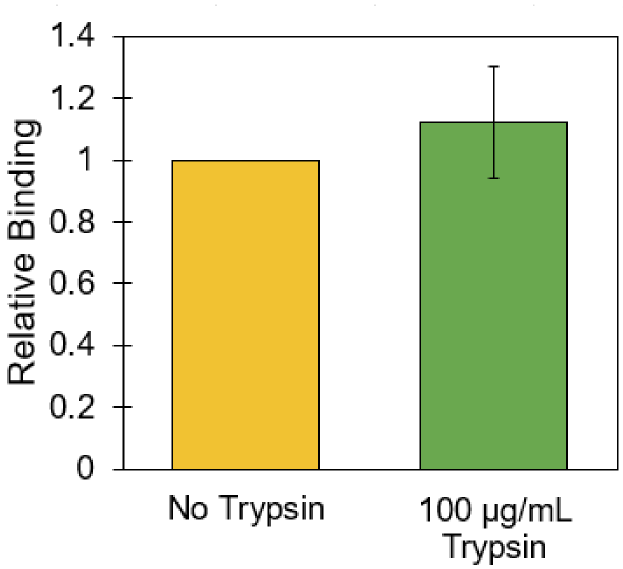

***Figure S7. SeV binding to SLBs is unaffected by trypsin treatment.*** *Single-virus binding measurements to supported lipid bilayers were performed with SeV with that had been untreated or treated with 100 µg/mL TPCK-trypsin. Binding is calculated relative to no trypsin treatment, and error bars are ± standard error of ≥3 sample replicates, and ≥10 separate image locations within each sample, with a propagated relative error as described previously* (2). *Lipid composition of target SLB = 2% GQ1b, 20% DOPE, 10% cholesterol, 0.05% Oregon Green-DHPE, and 67.95% POPC.*

<u>**S.6: Chemical structures of gangliosides GQ1b and GD1a**</u>

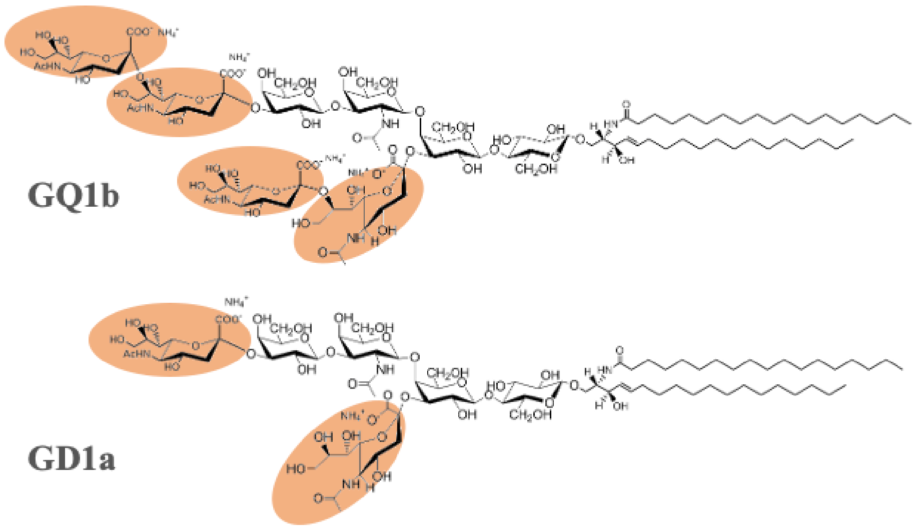

***Figure S8. Chemical structures of gangliosides GQ1b and GD1a, used as SeV receptors in our single virus lipid mixing experiments.*** *Sialic acid residues are highlighted in orange.*

### S.7: Supplementary References
